# Diffusion of lipids and GPI-anchored proteins in actin-free plasma membrane vesicles measured by STED-FCS

**DOI:** 10.1101/076109

**Authors:** Falk Schneider, Mathias P. Clausen, Dominic Waithe, Thomas Koller, Gunes Ozhan, Christian Eggeling, Erdinc Sezgin

**Affiliations:** MRC Human Immunology Unit, Weatherall Institute of Molecular Medicine, University of Oxford, OX39DS, Oxford, United Kingdom; MEMPHYS – Center for Biomembrane Physics, Department of Physics, Chemistry, and Pharmacy, University of Southern Denmark, Campusvej 55, 5230 Odense M, Denmark; Wolfson Imaging Centre Oxford, Weatherall Institute of Molecular Medicine, University of Oxford,OX39DS, Oxford, United Kingdom; Izmir International Biomedicine and Genome Institute (iBG-izmir), Dokuz Eylul University, Inciralti-Balcova, 35340 Izmir, Turkey; Department of Medical Biology and Genetics, Dokuz Eylul University Medical School, Inciralti-Balcova, 35340 Izmir, Turkey.

**Keywords:** plasma membrane biophysics, fluorescence correlation spectroscopy, lipid raft, GPI anchored proteins, STED microscopy, STED-FCS, giant plasma membrane vesicles

## Abstract

Diffusion and interaction dynamics of molecules at the plasma membrane play an important role in cellular signalling. These have been suggested to be strongly associated with the actin cytoskeleton. Here, we utilise super-resolution STED microscopy combined with fluorescence correlation spectroscopy (STED-FCS) to access the sub-diffraction diffusion regime of different fluorescent lipid analogues and GPI-anchored proteins (GPI-APs) in the cellular plasma membrane, and compare it to the diffusion regime of these molecules in cell-derived actin-free giant plasma membrane vesicles (GPMVs). We show that phospholipids and sphingomyelin, which undergo hindered diffusion in the live cell membrane, diffuse freely in the GPMVs. In contrast to sphingomyelin, which is transiently trapped on molecular-scale complexes in intact cells, diffusion of the ganglioside lipid GM1 suggests transient incorporation into nanodomains, which is less influenced by the actin cortex. Finally, our data on GPI-APs indicate two molecular pools in living cells, one pool showing high mobility with trapped and compartmentalized diffusion, and the other forming immobile clusters both of which disappear in GPMVs. Our data underlines the crucial role of the actin cortex in maintaining hindered diffusion modes of most but not all membrane molecules.

GPMVs: giant plasma membrane vesicles
GUVs: giant unilamellar vesicles
GPI-AP: Glycophosphatidylinositol anchored protein
PL: phospholipid
STED: Stimulated emission depletion
FCS: fluorescence correlation spectroscopy

## Introduction

The cellular plasma membrane is considered as a heterogeneous structure composed of various types of lipids and proteins, and this heterogeneity plays crucial roles in cellular signalling (Simons and Gerl, 2010). The underlying physical-chemical principles have been extensively studied for many years (Lingwood and Simons, 2010). A first comprehensive concept to describe the membrane structure and dynamics was the fluid mosaic model. This model suggested a homogenous multi-component system wherein, on the long-range, processes are based on free Brownian motion, yet on a short-range, interactions of lipids and proteins can form small-scale heterogeneous complexes (Singer and Nicolson, 1972). Later models suggested a more elaborated sub-organisation of the membrane into functional domains (Simons and Ikonen, 1997). These nano-domains, referred to as membrane rafts, were proposed to be enriched in cholesterol and saturated lipids (Simons and Ikonen, 1997) and to be highly dynamic (Pike, 2006; Sezgin *et al.*, 2015). Besides the relatively general raft concept, nano-clusters of specific membrane components have been reported. GPI-anchored proteins (Varma and Mayor, 1998), ganglioside GM1 (Yuan and Johnston, 2001), sphingomyelin (Guyomarc'h *et al.*, 2014), or specific immune receptor clusters (Dustin and Groves, 2012) are only a few examples that were shown to build up nano-scale heterogeneous structures at cellular membranes (Saka *et al.*, 2014).

Although the existence of rafts as a general organising concept of the plasma membrane remains under debate (Klotzsch and Schuetz, 2013), there is a consensus on the presence of membrane heterogeneity in terms of structure and dynamics (Sezgin *et al.*, 2015). The latter, for instance, is often quantitatively investigated by measuring diffusion of proteins and lipids in the cellular plasma membrane. Such diffusion measurements were used to determine the molecular mobility in segregated domains (Kahya *et al.*, 2003; Sezgin and Schwille, 2012), to elucidate the binding dynamics of cell surface receptors (Yu *et al.*, 2009), to investigate the influence of the underlying cytoskeleton structure on membrane dynamics (Kusumi *et al.*, 2005; Kusumi *et al.*, 2010; Mueller *et al.*, 2011; Andrade *et al.*, 2015; Fujiwara *et al.*, 2016; Koster *et al.*, 2016; Koster and Mayor, 2016), and the formation of transient interactions (Eggeling *et al.*, 2009; Honigmann *et al.*, 2014), to name a few.

An important tool to measure molecular mobility in membranes is fluorescence correlation spectroscopy (FCS) (Fahey *et al.*, 1977; Schwille *et al.*, 1999). In FCS, an apparent diffusion coefficient *D* is determined from the average transit time of fluorescently tagged molecules moving into and out of a microscope’s observation spot. FCS data recorded for different observation spot sizes confirmed that different components of the plasma membrane diffuse not only with different velocities but also with different diffusion modes (Wawrezinieck *et al.*, 2005; Eggeling *et al.*, 2009). The diffusion mode defines how the apparent diffusion coefficient of molecules changes with the size of the observation spot (Eggeling, 2015). An illustration of proposed diffusion modes and their possible underlying mechanisms are shown in Figure 1. For a molecule undergoing free (Brownian) diffusion the apparent diffusion coefficient is not dependent on the size of the observation spot (Fig 1A). This is different for hindered diffusion. For example, *D* will decrease with decreasing observation spot size diameter *d*, if molecules are transiently immobilized or trapped in e.g. immobilized or slow-moving molecular complexes (Figure 1B) (Eggeling *et al.*, 2009; Mueller *et al.*, 2011). On the other hand, *D* will increase towards smaller spot size *d* when molecules undergo hop or compartmentalized diffusion (Figure 1C) (Fujiwara *et al.*, 2002; Clausen and Lagerholm, 2013; Andrade *et al.*, 2015). In case molecules are transiently incorporated in domains, where their diffusion is slowed down, *D* will decrease towards smaller *d* in a similar manner as for the trapped diffusion, but will level out or slightly increase again approximately at the diameter of the domain size (Figure 1D) (Honigmann *et al.*, 2013; Guzman *et al.*, 2014; Sachl *et al.*, 2016). Yet, this levelling out at lower *d* can well be obtained when both significant hop and trapped diffusion is present simultaneously (Mueller, 2012). As outlined, the *D(d)* dependence is usually determined by measuring FCS data for different observation sizes (spot-variation FCS) (He and Marguet, 2011). This can either be done on conventional confocal microscopes with diffraction-limited observation spots *d* > 200 nm (Wawrezinieck *et al.*, 2005), or on a super-resolution STED microscope with *d* < 200 nm (Eggeling *et al.*, 2009). The latter has the advantage that diffusion is measured closer to the length scales of the features causing hindrances in diffusion.

**Figure 1.**
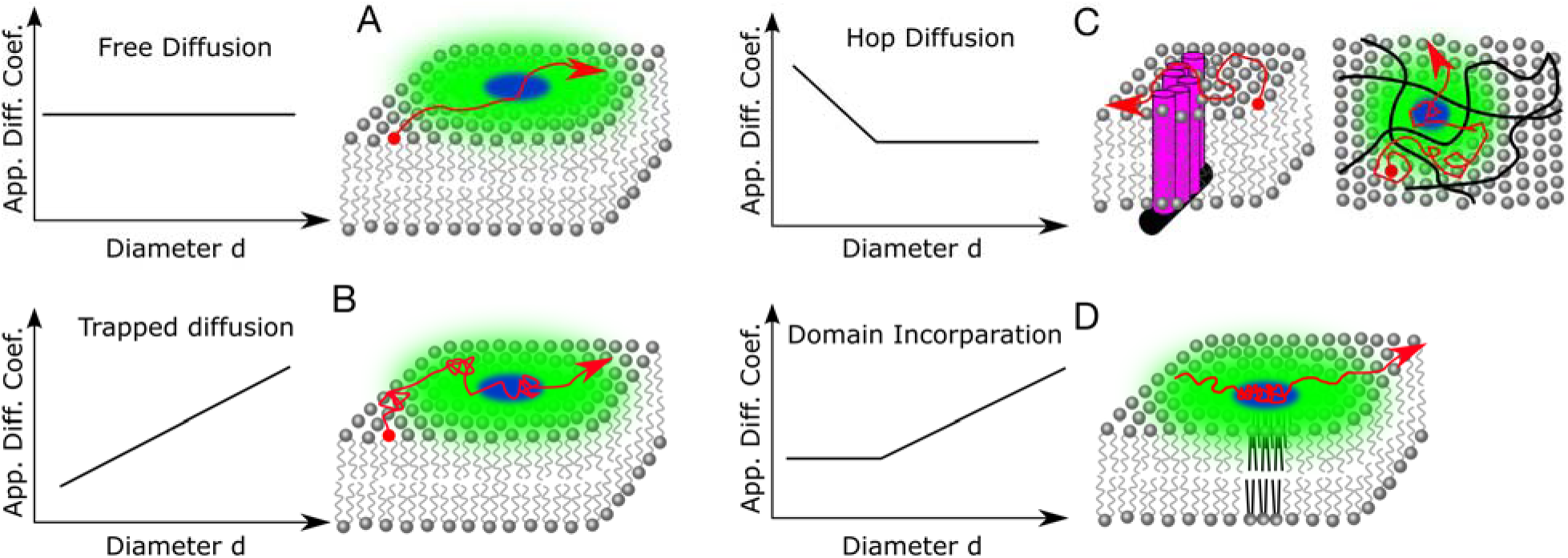
Sketch of possible diffusion modes in the cellular plasma membrane revealed by determination of the apparent diffusion coefficient *D* (App. Diff. Coef.) for different observation spot diameters *d* (left panels), and the potential molecular mechanisms behind these diffusion modes (right panels; red: molecular diffusion track, green and blue: large confocal and small STED microscope observation spots respectively, grey: lipids, black: actin cytoskeleton, purple: actin-anchored transmembrane proteins). A) Free diffusion; *D* remains constant. B) Trapped diffusion; *D* drops with decreasing *d*, presumably due to transient binding to immobile or slow-moving interaction partners, possibly assisted by the actin cytoskeleton. C) Hop diffusion; *D* increases towards small *d* due to compartmentalization of the membrane by the cortical actin meshwork and transmembrane proteins associated with it, leading to fast diffusion insight the compartments as probed at small *d* and hindrance in crossing from one to the next compartment as observed at large *d*. D) Transient domain incorporation; *D* decreases towards smaller *d* in a similar manner as in the trapping diffusion, but levels out or slightly increases again approximately at the diameter of the domain size, since diffusion is slowed down inside the domains, e.g. due to an increased molecular order.

Unfortunately, the exact mechanisms underlying the diffusion modes could not yet be unequivocally resolved. For example, trapped diffusion is thought to be caused by transient interactions with extremely slow or immobile membrane components, assisted by cholesterol and the actin cytoskeleton (Figure 1B) (Eggeling *et al.*, 2009; Mueller *et al.*, 2011). Hop diffusion is proposed to be induced by confinement from the cortical actin cytoskeleton meshwork and transmembrane proteins associated with it (Figure 1C) (Ritchie *et al.*, 2003; Kusumi *et al.*, 2010; Andrade *et al.*, 2015). The domains to which molecules may transiently incorporate may follow the lipid raft idea (Wawrezinieck *et al.*, 2005). Many of the above indications have been obtained from the experiments using perturbing drugs, such as cholesterol oxidising drugs or those de-polymerising the actin cytoskeleton. Yet, the action of these drugs might be manifold. Therefore, experiments under more controlled conditions and in minimal systems containing only the essential elements are necessary to confirm the molecular mechanisms inducing the different hindrances in diffusion.

In this study, we use STED-FCS to compare the diffusion modes of fluorescently labelled lipids and GPI-anchored proteins in the plasma membrane of live cells and cell-derived membrane vesicles, so called giant plasma membrane vesicles (GPMVs). GPMVs display an excellent model of the cellular membrane (Scott and Maercklein, 1979) since they contain most of the natural membrane components but lack the actin cytoskeleton (Baumgart *et al.*, 2007). Our data underlines the crucial role of the actin cortex in maintaining hindered diffusion modes of most but not all membrane molecules, since hindered diffusion is to a large extend abolished in actin-free GPMVs.

## Results and Discussion

### Giant Plasma Membrane Vesicles lack organized cytoskeleton

We first tested the distribution of actin in the GPMVs to confirm the lack of actin cytoskeleton in these vesicles. We visualized the actin organization in adherent and suspended cells and in giant plasma membrane vesicles (GPMVs) derived thereof. Figure 2A shows the filamentous actin (F-actin) organisation in live adherent CHO cells labelled with Lifeact-GFP where cortical actin cytoskeleton clearly becomes visible as a bright structure beneath the plasma membrane. In contrast, in CHO cells-derived GPMVs, we observed no such cortical actin network underneath the membrane; instead actin was homogenously distributed inside the GPMVs (Figure 2A). Similarly, monomeric actin (or globular actin, G-actin, labelled with the fluorescent protein Citrine) showed an increased preference to the cellular cortex beneath the plasma membrane of suspended Jurkat T cells, while it is homogeneously distributed throughout T cell derived GPMVs (Figure 2B). With this data, we confirm that the cortical actin cytoskeleton is abolished in GPMVs.

**Figure 2.**
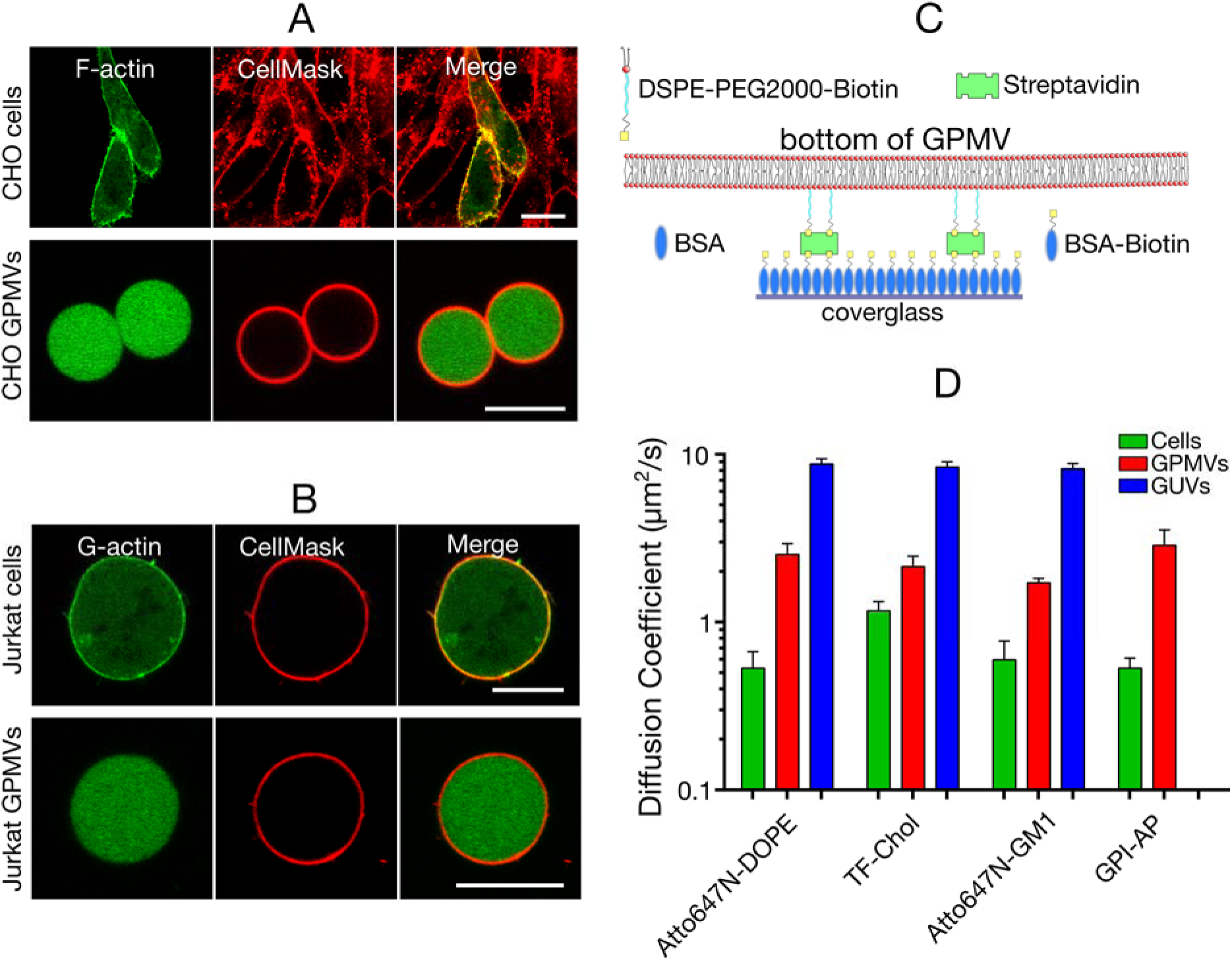
Distribution of actin and diffusion of lipids and GPI-AP in intact cells and GPMVs. A, B) Confocal images of the equatorial plane of A) adherent CHO cells (upper panels) and GPMVs derived thereof (lower panels) expressing Lifeact-GFP F-actin (green), and of B) suspended Jurkat T-cells (upper panels) and GPMVs derived thereof expressing fluorescent G-actin (green). In both cases the plasma membrane is visualized with CellMask (red). Scale bars: 10 µm.C) Sketch of the immobilization strategy for GPMVs. D) Diffusion coefficients obtained by confocal FCS recordings of Atto647N-DOPE, Topfluor-cholesterol (TF-chol), Atto647N-GM1 and GPI-AP (labelled via a SNAP-tag) in live CHO cells (green), GPMVs (red), and GUVs (blue). Error bars are standard deviations of the mean of at least 10 measurements on different cells or vesicles.

### Diffusion in GPMVs is faster than in cells, but slower than in GUVs

Cortical actin has tremendous impact on the membrane structure and dynamics which was elucidated using actin-targeting drugs (Kusumi *et al.*, 2010; Koster *et al.*, 2016; Koster and Mayor, 2016; Saha *et al.*, 2016). In order to understand the impact of complete absence of actin cytoskeleton on molecular diffusion in membranes, we tested the diffusion of fluorescent (Atto647N- or TopFluor-labelled) analogues of the phospholipid 1,2-dioleoyl-sn-glycero-3-phosphoethanolamine (Atto647N-DOPE), cholesterol (TF-Chol) and the glycolipid GM1 (Atto647N-GM1) in the plasma membrane of live CHO cells, in GPMVs derived thereof, and in artificial free-standing giant unilamellar vesicles (GUVs; 100% 1,2-dioleoyl-sn-glycero-3-phosphocholine, DOPC) using confocal fluorescence correlation spectroscopy (FCS). FCS measurements in the adherent cells and GUVs were straightforward, since both settled down efficiently on the microscope cover glass and remained immobile for long enough time. In contrast, almost all GPMVs were still mobile, making the 10-30 s long FCS measurements challenging. Therefore, we immobilised GPMVs by incorporating a small amount of PEGylated- and biotinylated lipid into the membrane of GPMVs (see methods for details) and depositing them on streptavidin functionalized microscope cover glass surface (Figure 2C). We first confirmed that immobilisation did not influence the diffusion in the membrane of GPMVs by comparing FCS data obtained at the basal and apical membrane of immobilized GPMVs (Figure S1A) and in rarely appearing, non-moving non-immobilised GPMVs (Figure S1B, C). In the following, we solely determined molecular diffusion at the basal membrane of immobilized GPMVs, since this was the least challenging and most reproducible measurement mode.

Diffusion of all tested molecules was significantly faster in GPMVs than in living cells but still significantly slower than in the GUVs (Figure 2D). We account these differences due to the missing hindrance from the actin cytoskeleton in GPMVs (also suggested by previous measurements (Fujiwara *et al.*, 2016)) and less molecular crowding in GUVs (Houser *et al.*, 2016), respectively. All tested molecules diffused with similar diffusion coefficients in GUVs (D ≈ 8.5 µm^2^/s). In intact cells, Atto647N-DOPE and Atto647N-GM1 had similar diffusion coefficients (D ≈ 0.5 µm^2^/s), while TF-Chol was notably faster (D ≈ 1.2 µm^2^/s, in accordance with previous findings (Solanko *et al.*, 2013; Hiramoto-Yamaki *et al.*, 2014)). In GPMVs, Atto647N-DOPE and TF-Chol had similar diffusion coefficients (D ≈ 2.5 µm^2^/s). Larger increase in mobility for Atto647N-DOPE compared to TF-Chol suggests a stronger confinement of phospholipids by the cortical actin cytoskeleton than of cholesterol. In comparison, Atto647N-GM1 diffused rather slowly in GPMVs (D ≈ 1.5 µm^2^/s), which highlights a hindrance in diffusion not associated with the actin cortex. We also tested the diffusion of a fluorescently tagged GPI-anchored protein (GPI-AP), which was similar to that of Atto647N-DOPE both in cells and GPMVs.

### Hindered diffusion is abolished in GPMVs

Hindered diffusion in intact living cells has been reported several times for sphingolipids and phospholipids, specifically trapped diffusion in the case of sphingomyelin or GM1 (Eggeling *et al.*, 2009; Mueller *et al.*, 2011; Sezgin *et al.*, 2012b) and hop diffusion in the case of phospholipids such as DOPE and DPPE (Fujiwara *et al.*, 2002; Clausen and Lagerholm, 2013; Andrade *et al.*, 2015). We investigated whether these hindered diffusion modes also appeared in GPMVs compared to intact PtK2 cells. We tested the diffusion modes of four different lipid analogues, all labelled with the organic dye Atto647N: a saturated (Atto647N-DPPE) and an unsaturated (Atto647N-DOPE) phospholipid as well as sphingomyelin (Atto647N-SM) and Atto647N-GM1. In intact cells, Atto647N-DPPE was diffusing freely (Figure 3A, B, as expected for PtK2 cells (Eggeling *et al.*, 2009)), while we found two pools of Atto647N-DOPE (Figure 3C, D). One pool showed slight hop diffusion (dashed green line P1 in Figure 3C), and the other free diffusion (solid green line P2 in Figure 3C). Atto647N-SM and Atto647N-GM1 showed trapped-like diffusion in cells (Figure 3E-H). In contrast, diffusion of DOPE, DPPE and SM was free in GPMVs with an up to 5-fold increased mobility (D ≈ 2.5 µm^2^/s, red lines in Figure 3A, C, E). The abolishment in hindered diffusion in GPMVs is highlighted further by plotting the ratio D_STED_/D_Conf_ of the apparent diffusion coefficients determined from the FCS data of the STED (D_STED_) and confocal (D_Conf_) recordings. D_STED_/D_Conf_ = 1 depicts free, D_STED_/D_Conf_

>1 hop and D_STED_/D_Conf_ <1 trapped diffusion. In intact cells, we found the following values: D_STED_/D_Conf_ ≈ 1 for Atto647N-DPPE (Figure 3B) as well as for one pool of Atto647N-DOPE (Figure 3D), D_STED_/D_Conf_ ≈ 1.8 for the other pool of Atto647N-DOPE (Figure 3D), D_STED_/D_Conf_ ≈ 0.38 for Atto647N-SM (Figure 3F) and D_STED_/D_Conf_ ≈ 0.55 for Atto647N-GM1 (Figure 3H). These values increased for DOPE, DPPE and SM in GPMVs, reaching values close to D_STED_/D_Conf_ ≈ 1. The most obvious reason for the disappearance of hindered diffusion in GPMVs is the involvement of the cortical actin cytoskeleton in these hindrances, delivering compartmental confinements for hop diffusion as well as anchoring sites for immobilization of molecules that potentially serve as binding partners for transient trapping.

**Figure 3.**
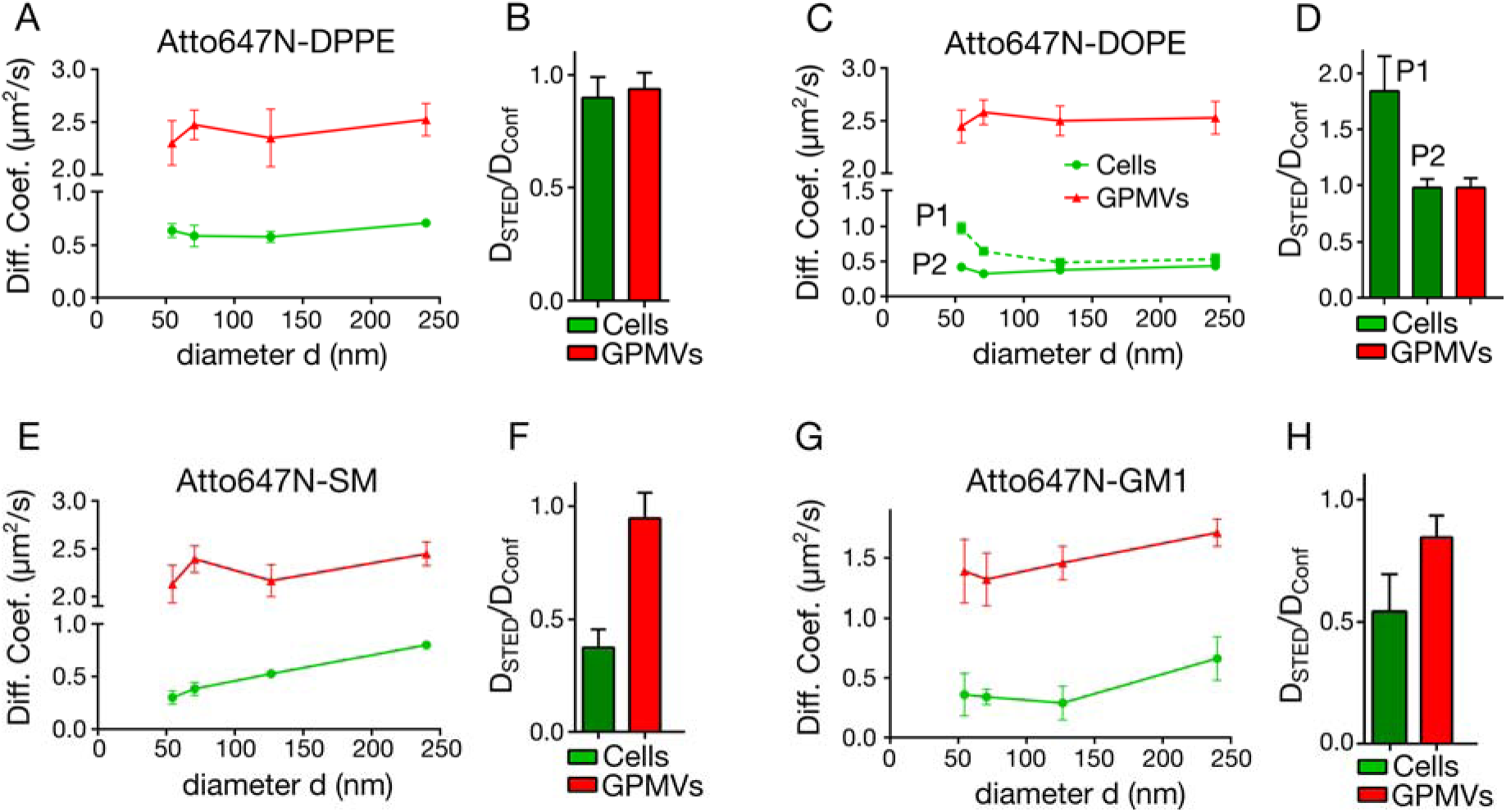
Diffusion modes of Atto647N-labeled lipid analogues in intact cells (green) and GPMVs (red) revealed by STED-FCS. A, C, E, G) Dependencies of values of the apparent diffusion coefficients *D* (Diff. Coef.) on the diameter *d* of the observation spot, and B, D, F, H) ratios D_STED_/D_Conf_ taken from these *D(d)* dependencies (with D_STED_ and D_Conf_ values of *D* at 60 nm and 240 nm, respectively) for A, B) Atto647N-DPPE, C, D) Atto647N-DOPE showing two different pools P1 and P2, E, F) Atto647N-SM, and G, H) Atto647N-GM1. Error bars are standard deviations of the mean of at least 10 measurements on different cells or vesicles.

A closer look on the *D(d)* dependency of Atto647N-GM1 suggests that this lipid transiently incorporates into domains of ≈120 nm in diameter in cells, instead of pure trapped diffusion as for Atto647N-SM (compare Figure 3E and G). As mentioned earlier, in purely trapped diffusion the molecule is not moving during the transient halt, while during domain incorporation the “trapped” molecule is still diffusing within the borders of domain with relatively slower velocity (see Figure 1B and D). Further, as already highlighted in Figure 2, the increase in mobility of Atto647N-GM1 between intact cells and GPMVs is about 3-fold, significantly lower than for the other lipids in GPMVs (≈5-fold, Figure 2D). This relatively slower diffusion results from still hindered diffusion of Atto647N-GM1 in GPMVs as indicated clearly by D_STED_/D_Conf_ ≈0.83 (Figure 3H). The corresponding diffusion mode (Figure 3G) suggests residual transient incorporation into nanodomains. In Figure 1D, we pointed out that a *D(d)* dependence similar to transient domain incorporation can well result from the presence of both trapped and hop diffusion. Yet, hop or compartmentalized diffusion has so far only been observed in the presence of a strong cortical actin meshwork. This favours the model of transient domain incorporation of GM1 in GPMVs which was suggested for artificial vesicles previously (Lozano *et al.*, 2013; Amaro *et al.*, 2016). Thus, our data indicates that the actin cytoskeleton influences diffusion of the phospholipids and sphingomyelin more significantly than of GM1.

### Diffusion of GPI-anchored proteins in cells and GPMVs

Next, we investigated the diffusion mode of GPI-anchored proteins in intact cells and GPMVs. We expressed a GPI anchored SNAP-tag domain (GPI-SNAP) in PtK2 cells and labelled it with the dye Abberior Star Red. STED-FCS data suggested transient incorporation into domains of roughly 75 nm in diameter (Figure 4A). As for most of the lipid analogues, hindered diffusion was abolished in GPMVs (D_STED_/D_Conf_ ≈ 1, Figure 4B) indicating the influence of the actin cytoskeleton on the diffusion dynamics of GPI-APs. In the case of the GPMV measurements, the GPI-SNAP could not be labelled efficiently enough, yielding extremely low signal. We therefore used RFP (red fluorescent protein) tagged GPI-APs (GPI-RFP) whose fluorescence signal was further amplified using an Atto647N-labelled nanobody against RFP.

**Figure 4.**
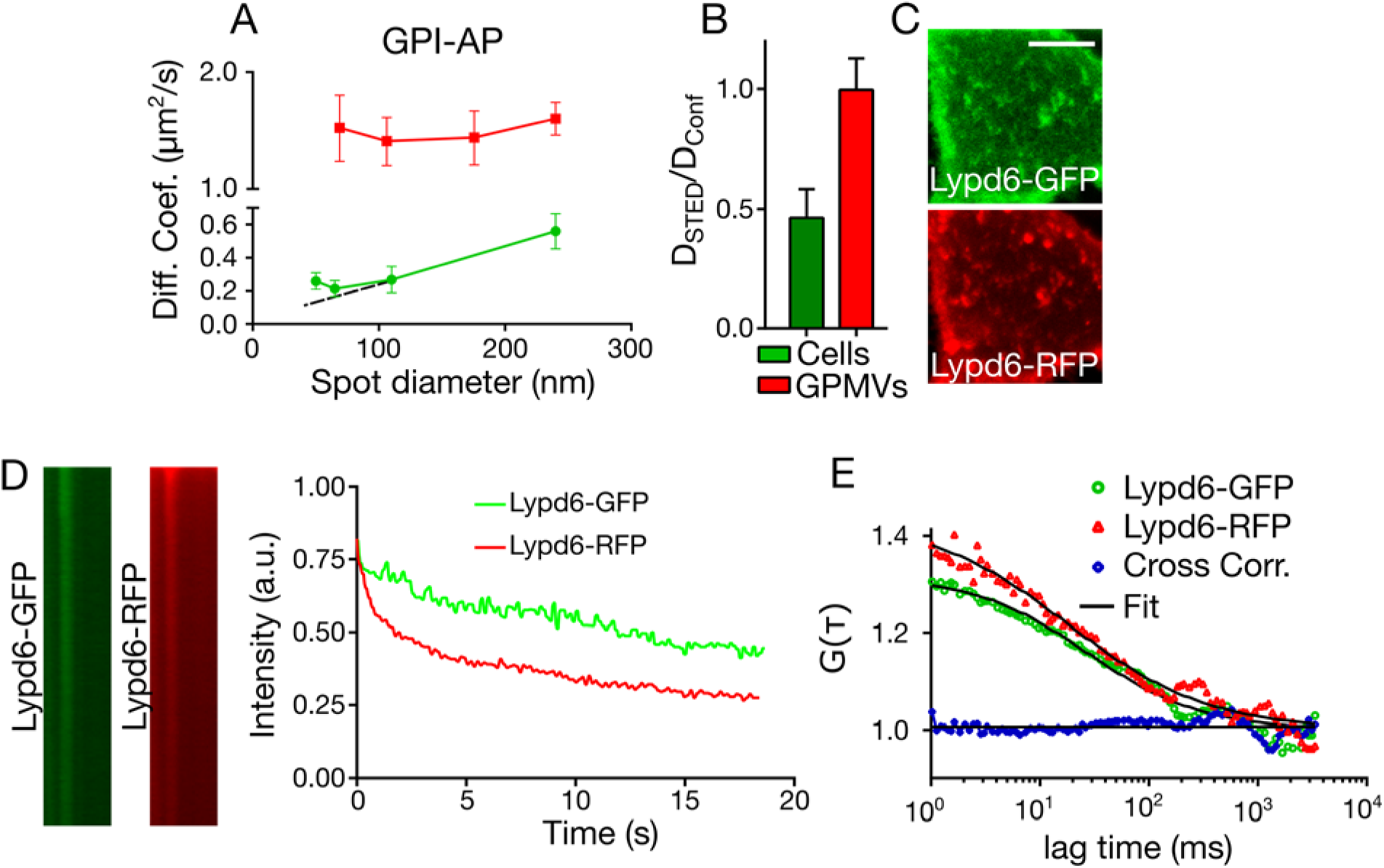
Diffusion of GPI-AP in cells and GPMVs. A) Dependence of the apparent diffusion coefficient *D* on the diameter *d* of the observation spot for GPI-SNAP in PtK2 cells (red, showing deviation from pure trapped diffusion, dashed black line) and GPI-RFP labelled with a Atto647N-tagged nanobody in (red), and B) corresponding ratios D_STED_/D_Conf_ (with D_STED_ and D_Conf_ values of D at 60 nm and 240 nm, respectively). Error bars are standard deviations of the mean of at least 10 measurements on different cells or vesicles. C) Confocal images of cells expressing GPI-anchored Lypd6 proteins labelled with GFP (green, upper panel) and RFP (red, lower panel), showing overlapping bright clusters (Pearson Correlation Coefficient 0.51 ± 0.1). Scale bar 10 µm. D) Fluorescence signal over time from the Lypd6 clusters for the GFP (green) and RFP (red) label: (left panels) line-scans (space, x-axis) over time (y-axis from top to bottom) with a bright signal spot fading over time, and (right panel) corresponding fluorescence intensity over time. E) Auto-correlation (FCS, red and green) and cross-correlation (FCCS, blue) functions of Lypd6-GFP and Lypd6-RFP in the plasma membrane of live PtK2 cells, together with a fit to the data (black), disclosing high mobility of both Lypd6-GFP and Lypd6-RFP (FCS data) but no co-diffusion (FCCS data).

We next investigated the hindrance in diffusion of GPI-APs further. STED microscopy images of the spatial organization of the GPI-APs revealed no <60 nm domains, but rather ≈100 nm large, bright and sparse clusters (Figure S2). We explored these clusters further using a more biologically relevant GPI-AP, for which we expressed GFP-and RFP-tagged Lypd6 proteins in PtK2 cells (Lypd6-GFP and Lypd6-RFP, respectively). Lypd6 is a crucial GPI-AP involved in Wnt/β-catenin signalling (Oezhan *et al.*, 2013). Two-colour images of Lypd6-GFP and Lypd6-RFP confirmed the existence of the bright and sparse clusters, with Lypd6-GFP and Lypd6-RFP spots overlapping substantially in space (Pearson Correlation Coefficient 0.51 ± 0.1) (Figure 4C). Disappearance of the clusters in GPMVs demonstrates their maintenance is dependent on actin (Figure S3A). The clusters were also immobile or very slowly moving in cells (Movie S1). Further, repetitive scanning over these visible clusters showed quickly fading fluorescence signals for both GFP and RFP (Figure 4D), which reveals that there is no fast interchange or replenishment of GPI-APs within these clusters. Thus, these clusters cannot be involved in the diffusion dynamics of the GPI-APs that we have observed in the STED-FCS experiments. Yet, smaller nanodomains might exist into which GPI-APs transiently incorporate. These nanodomains might be invisible in the fluorescence images due to missing contrast between the incorporated and free-diffusing pool of GPI-APs (Eggeling *et al.*, 2009; Eggeling, 2015). Note that generally the GPI-APs tend to partition into more ordered lipid environments (so did Lypd6-GFP and Lypd6-RFP, as probed in phase-separated GPMVs, Figure S3B), supporting the lipid nanodomain or raft idea. To probe the existence of floating nanodomains that incorporate more than one GPI-AP at a time, we performed fluorescence cross correlation spectroscopy (FCCS) on the diffusion of Lypd6-GFP and Lypd6-RFP. If these proteins co-diffuse within a floating nanodomain, this should reveal itself by a non-zero amplitude in the FCCS data. Yet, we detected no noticeable cross correlation despite the mobility of both proteins (Figure 4E), contradicting the mobile GPI-AP nanodomain idea. Thus, the diffusion mode of the GPI-APs presumably results from a combination of trapped and hop diffusion, rather than transient incorporations into nanodomains.

Using STED-FCS, we have compared the diffusion modes of several fluorescent lipid analogues and GPI-APs in the plasma membrane of intact cells and cell-derived GPMVs. In all cases, diffusion was much faster in GPMVs and hindrances in diffusion weakened or mostly abolished. Fluorescence images of the cortical actin demonstrated the non-existence of this network in GPMVs, highlighting the essential role of the actin cortex in maintaining hindered diffusion modes. Specifically, DPPE, DOPE, and SM lipid analogues showed free, hop and trapped diffusion in intact PtK2 cells respectively, and they all exhibited free diffusion with 5-fold increased diffusion coefficients in GPMVs. In contrast, the observed diffusion characteristics of GM1 ganglioside lipids suggested transient incorporations into nanodomains both in intact PtK2 cells and GPMVs derived thereof. However, this diffusion mode was significantly less pronounced in the GPMVs. Nevertheless, the actin cortex seems to have much less influence on the diffusion dynamics of GM1 than, for example, of SM, confirming previous STED-FCS measurements using actin perturbing drugs (Mueller *et al.*, 2011). The distinct behaviour of the ganglioside GM1 might be crucial for its specific role in membrane bioactivity. GM1 is one of the most ubiquitously expressed glycolipids of the cell membrane and is essential in host-pathogen interactions (Basu and Mukhopadhyay, 2014). Further, GM1 was shown to form functional homo- (Amaro *et al.*, 2016) or hetero-domains with GPI-APs (Komura *et al.*, 2016).

Our data suggests that GPI-APs undergo both significant trapped and hop diffusion. Both hindrances were abolished in GPMVs, where we observed only free diffusion for GPI-APs. Further in cells, we identified a separate pool of GPI-APs that was immobile and formed bright, stable ≈100 nm large clusters. These clusters disappeared in GPMVs which suggests that they are dependent on an intact actin cytoskeleton. They are presumably those highlighted in previous studies (Raghupathy *et al.*, 2015), and different from the mobile pool. We found no sign of co-diffusing GPI-APs in the mobile pool, which may be the reason why previous studies have found that GPI-APs did not establish a raft-like membrane environment upon artificial cross-linking (Sevcsik *et al.*, 2015).

In conclusion, STED-FCS in combination with GPMVs serves as a powerful tool to reveal important details about molecular membrane organization and dynamics, particularly the role of organized cortical actin cytoskeleton. The ability to perform investigations on a plasma membrane-derived vesicles lacking the actin cytoskeleton as well as other non-equilibrium processes (such as endo- and exocytosis) presents a well-controlled experimental model system.

## Experiential Procedures

### Tissue culture and cell staining

PtK2 Cells were grown in DMEM (Sigma Aldrich) supplemented with 15 % FCS (Sigma Aldrich) and 1 % L-glutamine (Sigma Aldrich). CHO cells were cultured in DMEM/F12 (Lonza) supplemented with 10 % FCS and 1% L-Glutamine. Jurkat T cells were grown in RPMI 1640 (Sigma Aldrich) containing 10 % FCS, 1 % L-Glutamine and 10 mM HEPES. For GPMV production they were grown on 30 mm petri dishes and for diffusion measurements in the cellular membrane the cells were grown on 18 mm or 25 mm round cover slips (#1.5). Usually, the cells reached a confluency of 50-70 % before a measurement was performed. Cells were labelled in phenol-red free L15 medium (Sigma Aldrich) at a lipid concentration of 1 µg/mL (Atto647N-DPPE, -SM, -DOPE) for 15 min at room temperature. After washing twice with L15, measurements were performed immediately. The labelling with Atto647N-H-GM1 was performed in full medium (2 µg/mL) for at least 15 minutes at room temperature.

The lipid analogues Atto647N-DOPE, Atto647N-DPPE, Atto647N-SM were purchased from Atto-Tec. Atto647N-GM1 was synthesized by Prof. Guenter Schwartzmann (Bonn, Germany) (Eggeling *et al.*, 2009). TF-Chol is purchased from Avanti Polar Lipids. Transfections were performed using Lipofecatmine 3000 (Life Technologies) according to the manufacturer’s protocol. See ref (Oezhan *et al.*, 2013) for Lypd6-GFP, Lypd-RFP. GPI-SNAP was a gift from the lab of Prof. Stefan Hell (Goettingen, Germany).

SNAP labelling with Abberior Star Red (Abberior GmbH) was performed at 2 ug/mL in full medium at 37 °C for 30 minutes. After two washing steps with full medium at 37 °C (30 minutes each) STED-FCS measurements and imaging were performed in L15.

### Giant Plasma Membrane Vesicles (GPMVs)

GPMVs of PtK2 cells were prepared according to ref (Sezgin *et al.*, 2012a). Briefly, cells were grown to a confluency of approximately 70 %, washed with GPMV buffer (containing 10 mM HEPES, 150 mM NaCl, 2 mM CaCl_2_ pH 7.4), and after adding 2 mM DTT and 25 mM PFA the cells were incubated for at least 4 hours at 37 °C to allow to the PtK2 cells to produce a sufficient amount of GPMVs. The GPMVs containing supernatant was harvested. DSPE-PEG-biotin (Avanti Polar Lipids) was added to a final concentration of 2 ng/mL onto the GPMV suspension. After 1.5 hours GPMVs were labelled with the lipid analogues Atto647N-DPPE, Atto647N-SM and Atto647N-DOPE dissolved in ethanol. The fluorescent lipid analogues were added to the GPMV solution to a final concentration of 0.1 µg/mL, 0.2 µg/mL and 0.14 µg/mL, respectively. Atto647N-GM1 was dissolved in DMSO and was added to a final concentration of 4 µg/mL. After another 15 minutes of incubation the GPMVs were spun down at 10.000 rpm for 15 minutes, and 90 % of the supernatant replaced by fresh buffer. The last step was crucial for the removal of free biotinylated lipid. For the measurements of GPI-RFP an RFP-binding nanobody labelled with Atto647N (Chromotek), a kind gift from Dr. Christoffer Lagerhom, was added. The nanobody was diluted to 100 µg/mL in 4 % BSA (in PBS) and stored at 4 °C. For the labelling of GPMVs it was used at a final concentration of 2 µg/mL.

The GPMVs were immobilised using biotin and streptavidin. Glass cover slips were coated with a 5:1 mixture of BSA/biotinylated BSA (Sigma Aldrich) for 1.5 hours, extensively washed and incubated with a solution of 200 ng/mL streptavidin (Life Technologies) in PBS. After washing with GPMV buffer the biotinylated GPMVs were added. Measurements were performed after 20 minutes. Immobilised GPMVs were stable for several hours.

The formation of GPMVs from other cell lines (CHO or Jurkat) was much faster than for PtK2 (within 2 hours). Otherwise, these GPMVs were formed and treated the same way.

### Giant Unilamellar Vesicles (GUVs)

GUVs were prepared by electroformation (Garcia-Saez *et al.*, 2010). A solution of DOPC (Avanti Polar Lipids) with a concentration of 1 mg/mL in chloroform was spread on platinum wires. After solvent evaporation the electrodes were dipped into 300 mM sucrose. For 1 hour an electric field with a frequency of 10 Hz and a potential of 2 V, followed by a frequency of 2 Hz was applied. GUVs were handled with cut tips and measurements performed in PBS. Cover slips were coated with BSA. GUVs were labelled by adding the lipid analogues Topfluor-Chol (Avanti Polar Lipids), Atto47N-DOPE, Atto647N-SM or Atto647N-DPPE to a final concentration of 0.05 µg/mL. Atto647N-GM1 was used at a final concentration of 0.005 µg/mL.

### Supported Lipid Bilayers (SLBs)

Supported lipid bilayers were prepared by spin coating (Clausen *et al.*, 2015). The cover slips were previously cleaned and edged by piranha acid (3:1 sulfuric acid and hydrogen peroxide). Cover slips were stored in water for not longer than a week.

A solution of 1 mg/mL of DOPC in chloroform/methanol was spin coated on to a cover slip at 3,200 rpm for 45 seconds. The lipid bilayer formed by rehydration with SLB buffer containing 10 mM HEPES and 150 mM NaCl pH 7.4. The SLB was labelled with AbberiorStar Red-DPPE (Abberior GmbH) (approximately 1:2000 molar ratio) contained in the initial DOPC solution.

### Data acquisition and evaluation

Confocal FCS, FCCS data and imaging data for fluorescent Lypd6 and the GPMV top versus bottom diffusion measurements were taken on a Zeiss780 LSM inverted confocal microscope equipped with a 40x C-Apochromat NA 1.2 W Corr FCS objective.

All STED-FCS data were taken on a customized Abberior STED microscope (Abberior Instruments) as previously described in ref (Clausen *et al.*, 2014; Galiani *et al.*, 2016). The microscope was equipped with a hardware correlator (Flex02-08D, correlator.com, operated by the company’s software). The dyes were excited using a 640 nm pulsed diode laser (PicoQuant, 80 MHz repetition rate) with an average excitation power of 5-10 µW at the objective (UPlanSApo 100x/1.4 oil, Olympus). For STED recordings, fluorescence emission was inhibited using a tuneable pulsed laser at 780 nm (Mai Tai, Newport; 80 MHz repetition rate). The microscope was operated using Abberior’s Imspector software. Fitting was performed with FoCuS-point fitting software as described in detail in ref (Waithe *et al.*, 2015).

The FCS data were fitted to a 2D diffusion model including one dark-state kinetics (with a relaxation time of 5 µs) for all SLB, GUV and GPMV measurements and an additional dark-state kinetics (with a relaxation time of 100 µs) for the data of all Atto647N-labelled lipids in the cellular membrane (as previously pointed out in (Mueller *et al.*, 2013)).

In the STED-FCS measurements, the diameter *d* of the observation spot is tuned by the average power *P*_*STED*_ of the STED laser. We performed STED-FCS measurements of AbberiorStar Red-DPPE in SLBs and GUVs at different *P*_*STED*_ to accurately calibrate the *d*(*P*_*STED*_) dependency. Since AbberiorStar Red-DPPE is diffusing freely in both model membranes, we can calculate the *d*(*P*_*STED*_) dependence using the following equation:

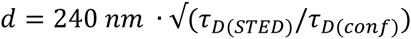

*τ*_*D(conf)*_ and *τ*_*D(STED)*_ are the transit times of the investigated molecules through the observation spot in confocal and at a certain STED laser power, respectively. The confocal observation spot diameter *d* = 240 nm was determined from confocal images of 20 nm Crimson beads (Life Technologies) spread out on a poly-L-Lysin (Sigma Aldrich) coated glass cover slip.

We employed calibration measurement performed on SLBs for the STED-FCS recordings on living cells, and those on GUVs for GPMV recordings.

The (apparent) diffusion coefficient *D* was calculated from *τ*_*D(conf)*_ and *τ*_*D(STED)*_ according to:

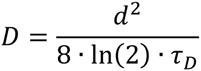

## Acknowledgments

We would like to thank Silvia Galiani for her continuous help on Abberior STED microscope, and Christoffer Lagerholm and Esther Garcia for help on the microscopes in the Wolfson Imaging Centre. E.S. is supported by EMBO long term (ALTF 636-2013) and Marie Skłodowska-Curie Intra-European Fellowships (MEMBRANE DYNAMICS 627088). This work is supported by the Wolfson Foundation (ref. 18272), the Medical Research Council (MRC, grant number MC_UU_12010/unit programmes G0902418 and MC_UU_12025), MRC/BBSRC/ESPRC (grant number MR/K01577X/1), and the Wellcome Trust (grant ref. 104924/14/Z/14).

